# *Mycobacterium tuberculosis* requires conditionally essential metabolic pathways for infection

**DOI:** 10.1101/2023.09.06.556627

**Authors:** Alisha M. Block, Parker C. Wiegert, Sarah B. Namugenyi, Anna D. Tischler

## Abstract

New drugs are needed to shorten and simplify treatment of tuberculosis caused by *Mycobacterium tuberculosis*. Metabolic pathways that *M. tuberculosis* requires for growth or survival during infection represent potential targets for anti-tubercular drug development. Genes and metabolic pathways essential for *M. tuberculosis* growth in standard laboratory culture conditions have been defined by genome-wide genetic screens. However, whether *M. tuberculosis* requires these essential genes during infection has not been comprehensively explored because mutant strains cannot be generated using standard methods. Here we show that *M. tuberculosis* requires functional phenylalanine (Phe) and *de novo* purine and thiamine biosynthetic pathways for mammalian infection. We used a defined collection of *M. tuberculosis* transposon (Tn) mutants in essential genes, which we generated using a custom nutrient-rich medium, and transposon sequencing (Tn-seq) to identify multiple central metabolic pathways required for fitness in a mouse infection model. We confirmed by individual retesting and complementation that mutations in *pheA* (Phe biosynthesis) or *purF* (purine and thiamine biosynthesis) cause death of *M. tuberculosis* in the absence of nutrient supplementation *in vitro* and strong attenuation in infected mice. Our findings show that Tn-seq with defined Tn mutant pools can be used to identify *M. tuberculosis* genes required during mouse lung infection. Our results also demonstrate that *M. tuberculosis* requires Phe and purine/thiamine biosynthesis for survival in the host, implicating these metabolic pathways as prime targets for the development of new antibiotics to combat tuberculosis.

**AUTHOR SUMMARY:** *Mycobacterium tuberculosis* causes more than 10 million new cases of active tuberculosis (TB) disease and 1.6 million deaths worldwide each year. Individuals with active TB must take a combination of four antibiotics for a minimum of 6-9 months to cure the infection. New anti-tubercular drugs are needed to simplify TB treatment and combat drug resistance. Here, we describe a novel collection of *M. tuberculosis* mutants lacking metabolic pathways essential for growth in standard laboratory conditions. Using these mutants, a mouse infection model, and deep sequencing we identified those metabolic pathways that *M. tuberculosis* also requires during infection. We find that *M. tuberculosis* mutants that cannot synthesize purine nucleotides, riboflavin, or certain amino acids are unable to grow in mice. We also find that mutant strains which cannot synthesize purine nucleotides or the amino acid phenylalanine die rapidly in laboratory cultures without nutrient supplementation, suggesting that new drugs targeting these pathways would kill *M. tuberculosis*. Overall, our work reveals multiple metabolic pathways that *M. tuberculosis* requires during infection, which could be pursued as new targets for development of anti-tubercular drugs.

## INTRODUCTION

*Mycobacterium tuberculosis* caused more deaths worldwide in 2021 than any other bacterial infectious agent, in part due to the complexity of tuberculosis (TB) treatment, which requires 6-9 months of therapy with multiple antibiotics [1, 2]. Multidrug-resistant *M. tuberculosis* (MDR-TB) accounts for ∼4.2% of TB infections and is more challenging to treat, requiring use of less effective second-line agents [1, 3]. Development of new anti-tubercular drugs will be critical to reduce the length of TB treatment and to combat antibiotic resistance. New drug regimens hold some promise for reducing TB treatment duration and for treating MDR-TB [4-6]. However, defining diverse new targets for TB drug development will be necessary to counteract antibiotic resistance, which has already been documented for all existing drugs and all new or repurposed TB drugs in the clinical pipeline [7]. Metabolic pathways that *M. tuberculosis* requires for *in vitro* growth and that are highly vulnerable to inhibition have been proposed as novel TB drug targets [8-11]. However, these metabolic pathways may not be required for *M. tuberculosis* growth during infection due to differences in nutrient availability in host tissues as compared to the *in vitro* growth medium. Defining the metabolic pathways that *M. tuberculosis* requires during mammalian infection will be essential to prioritize targets for TB drug development.

*M. tuberculosis* genes required during mammalian infection have been defined by genome-wide transposon (Tn) mutant screens in the mouse infection model [12-17]. However, these studies excluded analysis of genes that are essential for *M. tuberculosis* growth in standard culture conditions *in vitro* [11], as it is not possible to generate insertional mutations in these genes. In a few cases, auxotrophic *M. tuberculosis* mutant strains generated on medium supplemented with exogenous nutrients were used to demonstrate that specific central metabolic pathways are essential for growth in mice. These include pathways for synthesis of the amino acids Met, Arg, Pro and Trp [13, 18-21]. Other metabolic pathways such as coenzyme A, biotin, and trehalose synthesis were shown to be essential during mouse infection using *M. tuberculosis* conditional gene expression strains [22-24]. Each of these pathways are being pursued as potential targets for TB drug development.

However, not all metabolic pathways essential for *M. tuberculosis* growth *in vitro* are also required during mammalian infection. For example, the *nadABC* genes, which encode enzymes for *de novo* NAD+ synthesis, are essential *in vitro* but dispensable in the host [25, 26]. *M. tuberculosis* can scavenge the nicotinamide precursor from host tissue to synthesize NAD+ using an alternative salvage pathway [26]. This example highlights the importance of determining whether genes that *M. tuberculosis* requires for *in vitro* growth are also necessary during infection. Of the 625 genes annotated as essential for optimal growth of *M. tuberculosis* on standard Middlebrook medium [11], only a few have been directly tested to determine whether they are also required during infection because deletion mutant strains lacking these genes cannot be generated under standard culture conditions.

We previously developed an arrayed Tn mutant library using MtbYM rich medium, a custom medium that contains many additional carbon and nitrogen sources, amino acids, nucleotide precursors, co-factors and vitamins as compared to standard Middlebrook medium [27, 28]. Based on genome-wide transposon-sequencing (Tn-seq) analysis, 118 genes annotated as essential for optimal growth on Middlebrook 7H10 are non-essential on MtbYM rich [27]. These include genes required for synthesis of amino acids (Arg, Trp, Ile, Val, Leu, Glu, Pro, Ser, Tyr, Gln, Phe, Met, Cys), vitamins and co-factors (riboflavin, folates, pantothenate, NAD), and purine nucleotides, as well as certain catabolic pathways, including glycolysis [27]. We reasoned that our unique collection of Tn mutant strains selected on MtbYM rich medium could be exploited to rapidly assess whether *M. tuberculosis* requires these conditionally essential metabolic pathways for growth and survival during infection.

Here, we show that *M. tuberculosis* requires functional phenylalanine (Phe) biosynthesis and *de novo* purine nucleotide and thiamine biosynthesis pathways for growth in the mammalian host. We screened our collection of Tn mutants in conditionally essential genes in a mouse infection model using transposon sequencing (Tn-seq) and identified multiple central metabolic pathways that *M. tuberculosis* requires for fitness in host lung and spleen tissues. We confirm that mutation of *pheA* (Phe synthesis) or *purF* (purine/thiamine synthesis) causes death of *M. tuberculosis* in the absence of exogenous nutrient supplementation *in vitro* and in the lungs of aerosol-infected mice. Our results demonstrate roles for multiple metabolic pathways in *M. tuberculosis* fitness during infection and implicate the Phe and purine/thiamine biosynthetic pathways as novel targets for development of anti-tubercular drugs.

## RESULTS

*M. tuberculosis* genes that are essential for growth in standard Middlebrook medium have been defined by saturating Tn mutagenesis screens [11]. We previously constructed an arrayed Tn mutant library in the *M. tuberculosis* Erdman strain using MtbYM rich medium, which contains many additional carbon sources, amino acids, metabolic intermediates, vitamins and co-factors that are not present in Middlebrook medium [27, 28]. Our library contains 48 Tn mutants in genes annotated as “essential” in Middlebrook medium that are non-essential in MtbYM rich medium [11, 27, 28]. We refer to these genes as <u>M</u>iddlebrook-<u>Es</u>sential (M-ES) genes. To determine whether *M. tuberculosis* requires these M-ES genes for survival in the host, we isolated Tn mutants in M-ES genes from our arrayed library and screened them for fitness defects in mice using transposon sequencing (Tn-seq).

### Tn insertions in genes annotated as essential for growth on Middlebrook medium can be isolated on MtbYM rich medium

Our arrayed *M. tuberculosis* Erdman Tn library made on MtbYM rich medium contains a total of 105 unique Tn insertions in 48 M-ES genes (**S1 Table**). Of these, 20 mutants in 17 unique M-ES genes harbored the Tn insertion near the 5’ or 3’ end of the gene (within the first 5% or last 95% of the coding sequence). As these Tn insertions were unlikely to disrupt gene function, these mutants were excluded from this study (**S1C Table**, Tn mutants excluded). We recovered 45 of the Tn mutants in M-ES genes from their mapped locations in our arrayed library on MtbYM rich agar and confirmed the Tn insertion site by PCR (**S1A and S1B Table**).

We previously determined that some mutants in our MtbYM rich Tn library have defects in production of the outer membrane lipid phthiocerol dimycocerosate (PDIM) due to secondary mutations, unlinked to the Tn [28]. The M-ES Tn mutants might be PDIM deficient due to similar secondary mutations. As *M. tuberculosis* requires PDIM for virulence [17], we confirmed that the M-ES Tn mutants were PDIM-proficient prior to screening for fitness defects in mice. We used PCR and Sanger sequencing to test the M-ES Tn mutants for point mutations in PDIM biosynthesis genes that we identified in other Tn mutants from our library. This analysis identified 6 M-ES Tn mutants with mutations in genes required for PDIM production (**S1B Table**, unusable Tn mutants). We conducted whole genome sequencing on the remaining M-ES Tn mutants, which identified additional mutations predicted to prevent PDIM production in 17 Tn mutants and one mutant with multiple Tn insertions (**S1B Table**, unusable Tn mutants). These mutants were excluded from further analysis. We recovered Tn mutants in 19 M-ES genes which contained a single Tn insertion and no mutations in genes required for PDIM production, based on whole genome sequencing (**S1A Table**, Tn mutants in M-ES library). These M-ES Tn mutants, representing various metabolic pathways and essential processes (**Table 1**), were selected for analysis of growth *in vitro* and fitness defects in mice.

**Table 1.** Tn mutants in M-ES genes in the Tn library screened in mice.

| Gene | Annotation | Putative Function |
| --- | --- | --- |
| *purF* | amidophosphoribosyltransferase | *de novo* purine & thiamine biosynthesis |
| *purM* | phosphoribosyl-aminoimidazole synthase | *de novo* purine & thiamine synthesis |
| *purQ* | phosphoribosyl-formylglycinamide synthetase I | *de novo* purine & thiamine synthesis |
| *pheA* | prephenate dehydratase | Phe biosynthesis |
| *trpG* | anthranilate synthase component II | Trp biosynthesis |
| *trpB* | tryptophan synthase beta chain | Trp biosynthesis |
| *proC* | pyrroline-5-carboxylate reductase | Pro biosynthesis |
| *gltB* | ferredoxin-dependent glutamate synthetase | Glu biosynthesis |
| *ilvA* | threonine dehydratase | Ile biosynthesis |
| *panC* | pantoate-beta-alanine ligase | pantothenate biosynthesis |
| *ribA2* | GTP cyclohydrolase II | riboflavin biosynthesis |
| *ribG* | bifunctional riboflavin synthesis protein | riboflavin biosynthesis |
| *pabB* | *para*-aminobenzoic acid (PABA) synthase component I | PABA biosynthesis |
| *fhaA* | conserved protein with forkhead-associated domain | regulation of cell wall synthesis |
| *pstP* | serine-threonine phosphatase | regulation of cell wall synthesis |
| *ftsQ* | cell division protein | cell division |
| *ERDMAN_3321* | possible acyl transferase | unknown function |
| *ERDMAN_2739* | Obg GTPase | ribosome assembly |
| *ERDMAN_4254* | LigA DNA ligase | DNA damage repair |

### Tn mutants in central metabolic pathways have growth defects in Middlebrook 7H9 medium

To determine whether the M-ES genes disrupted by Tn insertion are required for *M. tuberculosis* growth in standard culture conditions, we conducted growth curves in Middlebrook 7H9 medium. Tn mutants in genes encoding various central metabolic pathway enzymes exhibited severe growth defects. These included Tn mutants in genes required for purine and thiamine biosynthesis (*purF*, *purM*, *purQ*; **Fig 1A**), biosynthesis of the amino acids Phe, Trp, Pro, Glu and Ile (*pheA*, *trpB*, *trpG*, *proC*, *gltB*, *ilvA*; **Figs 1B and 1C**), and riboflavin or *para*-amino benozoic acid (PABA) biosynthesis (*ribA2*, *ribG*, *pabB*; **Fig 1D**). The *panC*::Tn mutant was included in our study as a positive control, and as expected it also failed to grow in Middlebrook 7H9 (**Fig 1D**). *M. tuberculosis* requires PanC for synthesis of pantothenate (vitamin B5), an essential precursor in coenzyme A biosynthesis, in the absence of exogenous pantothenate [29]. These data demonstrate that each of these M-ES genes is essential for growth of *M. tuberculosis* Erdman in standard Middlebook medium, consistent with their annotation as essential for growth on Middlebrook 7H10 based on saturating Tn mutagenesis [11].

**Fig 1.**
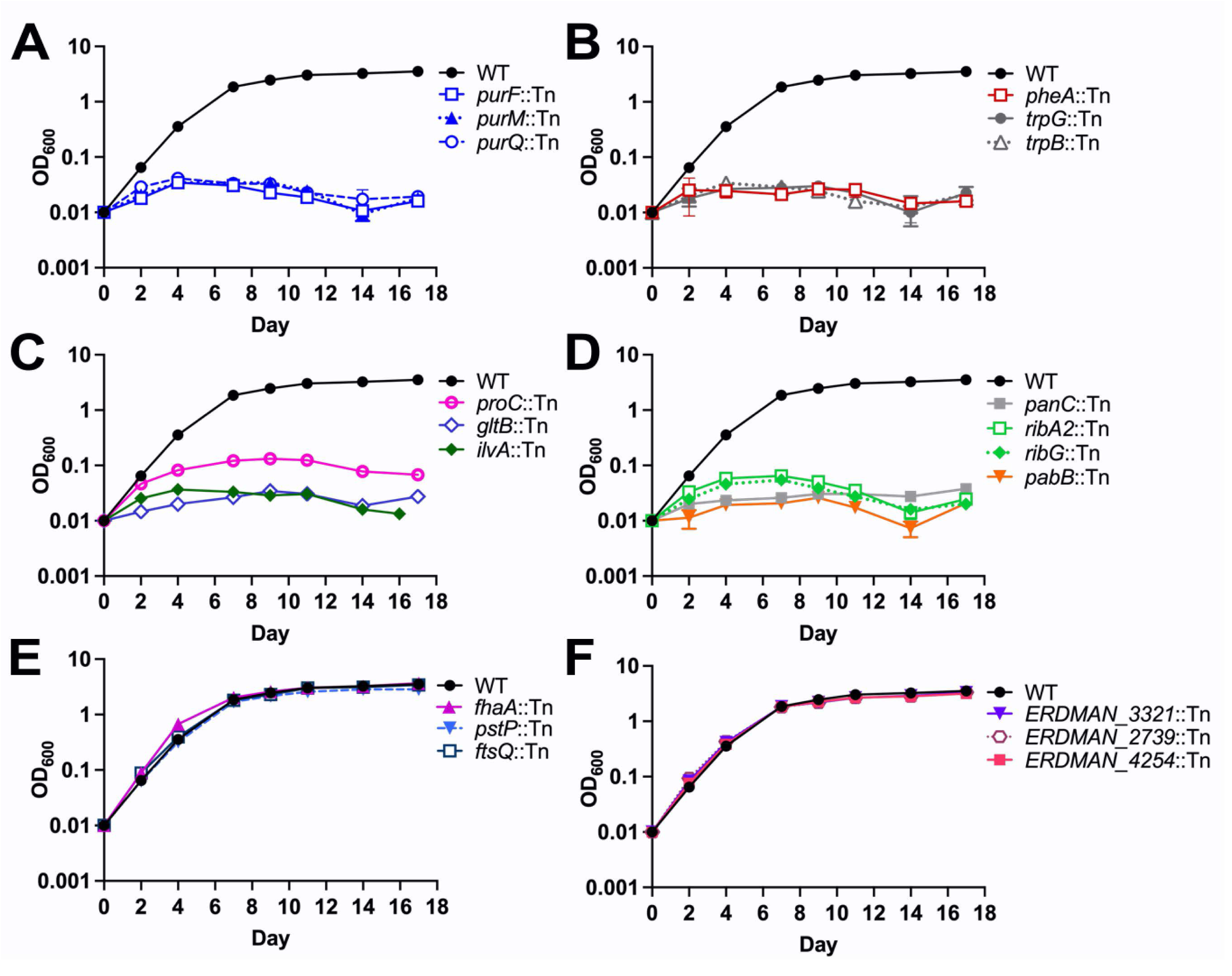
*M. tuberculosis* Tn mutants in genes encoding central metabolic enzymes exhibit growth defects in Middlebrook 7H9 medium. Wild-type *M. tuberculosis* Erdman and M-ES Tn mutant strains were grown in MtbYM rich medium, washed twice in PBS-T, and diluted to OD_600_ = 0.01 in Middlebrook 7H9 medium. Growth was monitored by measuring the OD_600_. Tn mutants are grouped according to predicted function: **(A)** purine and thiamine metabolism (*purF*, *purM*, *purQ*), **(B-C)** amino acid biosynthesis (*pheA*, *trpB*, *trpG*, *proC*, *gltB*, *ilvA*), **(D)** pantothenate, riboflavin, and *p-* amino benzoic acid (PABA) biosynthesis (*panC*, *ribA2*, *ribG*, *pabB)*, **(E-F)** Tn insertions within non-essential gene regions (*pstP, fhaA, ftsQ, ERDMAN_3321*) or misannotated genes (*ERDMAN_2739, ERDMAN_4254*). Data represent the mean ± standard error of three biological replicates.

Tn mutants in several other putative M-ES genes had no growth defects in 7H9 (**Figs 1E and 1F**). Four Tn mutants harbored insertions within non-essential domains of M-ES genes. PstP and FhaA regulate *M. tuberculosis* cell wall synthesis [30, 31] and have domains predicted to be non-essential by saturating Tn mutagenesis [11]. The *pstP*::Tn and *fhaA*::Tn mutants both harbor the Tn insertion in a non-essential region of the gene and grew normally (**Fig 1E**). *M. tuberculosis* requires the membrane protein FtsQ to regulate cell division [32]. Our *ftsQ*::Tn mutant harbors the Tn at 185 bp from the 5’ end, in a region predicted to be non-essential [11]. The *ftsQ*::Tn mutant exhibited no growth defect (**Fig 1E**), suggesting that truncated FtsQ lacking 62 amino acids of the N-terminal cytosolic domain can sustain normal cell division *in vitro.* Our data are consistent with the observation that the FtsQ periplasmic domain complements the cell division defects of an FtsQ-depleted strain [32]. *ERDMAN_3321* (*rv3034*), which encodes a putative acetyl transferase, is annotated as essential, but Tn insertions near the 5’ end of the gene confer a growth advantage [11]. The *ERDMAN_3321*::Tn mutant harbors the Tn insertion 151 bp from the 5’ end of the gene, within this non-essential region (**Fig 1F**).

We also isolated two mutants with Tn insertions in *M. tuberculosis* Erdman genes with annotations suggesting essentiality. *ERDMAN_2739* is annotated as *obg* [33]; in *M. tuberculosis* H37Rv *obgE* (r*v2440c*) encodes an essential GTPase that controls ribosome assembly [11, 34, 35]. *ERDMAN_4254* is annotated as a DNA ligase [33], with possible similarity to the essential DNA ligase *ligA* (*rv3014*) [11]. Whole genome sequencing revealed that the *ERDMAN_2739*::Tn insertion is within *pe_pgrs43* and the *ERDMAN_4254*::Tn insertion is within *espK*, both of which are non-essential genes [11]. Thus, it is not surprising that these mutants did not exhibit growth defects (**Fig 1F**). These data indicate that a subset of the Tn mutants we isolated do not disrupt functions essential for *M. tuberculosis* growth in standard Middlebrook medium.

### *M. tuberculosis* requires many central metabolic pathways for fitness in mice

To identify those M-ES Tn mutants with fitness defects in mice, we created and screened a library of Tn mutants in M-ES genes (**S1A Table**). We included in the M-ES Tn library five positive control Tn mutants in genes known to be required for *M. tuberculosis* fitness in mice: *panC*::Tn [29] and four independent Tn insertions in *phoP*, which encodes a response regulator essential for virulence [36, 37]. We also included five negative control Tn mutants in genes known to be dispensable for *M. tuberculosis* growth and survival in mice: *rpfA* and *rpfE* that encode redundant resuscitation promoting factors [38], *pstB* and *pstC1* that encode subunits of an alternate Pst phosphate transporter [39], and *tadA* that encodes a Flp pilin assembly ATPase [40]. Each of these negative control Tn mutants were predicted to be PDIM-proficient based on whole genome sequencing (**S1A Table**). The Tn mutants were grown individually in MtbYM rich liquid medium and equivalent amounts of each mutant were combined to generate the M-ES Tn library (**Fig 2A**). Bacteria in the M-ES Tn library were collected in triplicate for genomic DNA extraction as a control to confirm equal representation of each Tn mutant and aliquots of the library were frozen for use in experiments (**Fig 2A**).

**Fig 2.**
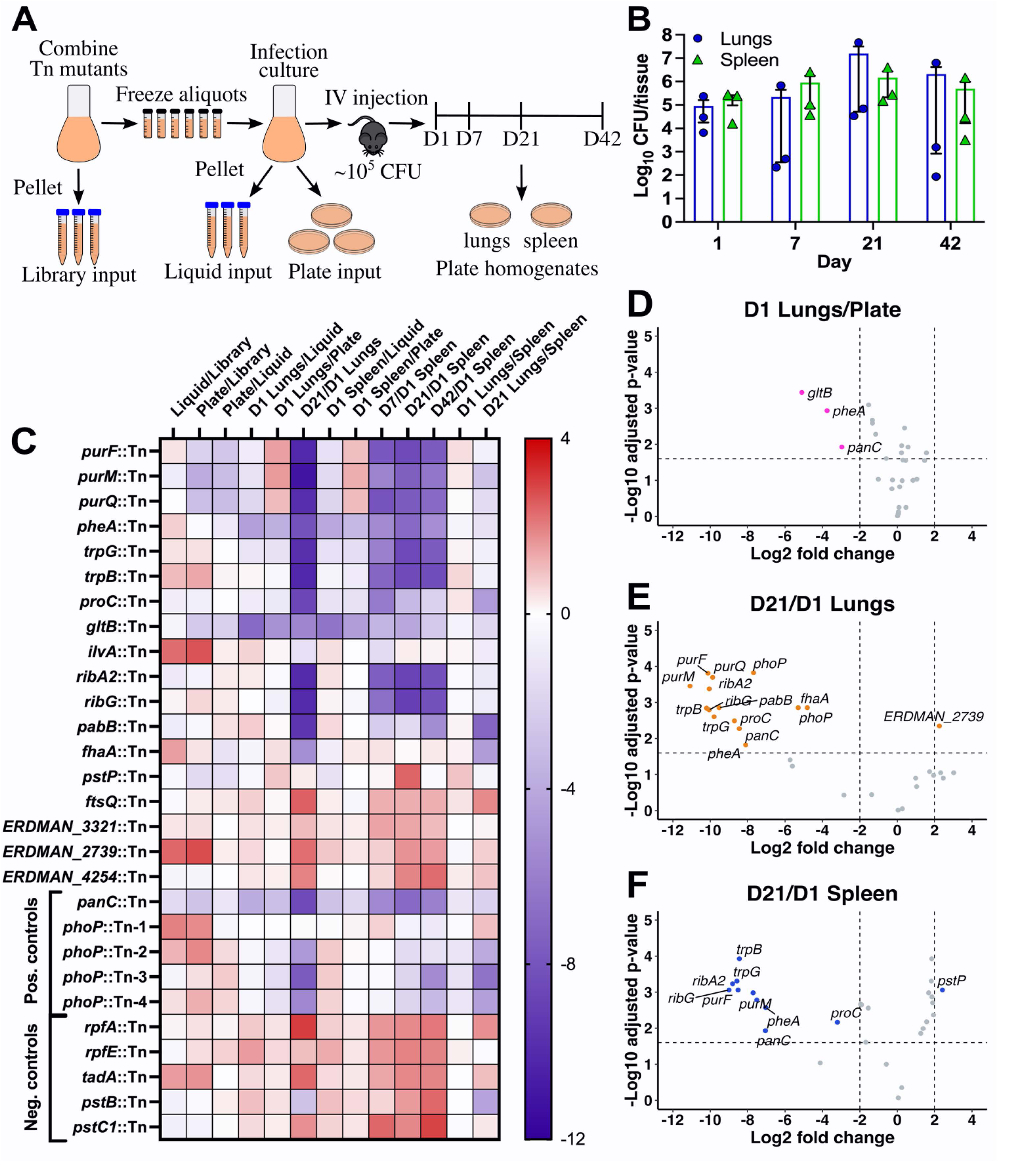
Tn-seq screen of the *M. tuberculosis* M-ES Tn library identifies multiple central metabolic pathways required for fitness in mice. **(A)** M-ES Tn-seq screen methods. Individual Tn mutants grown in MtbYM rich medium were mixed in equal abundance to create the M-ES Tn library. Triplicate M-ES Tn library samples were collected for Tn-seq (Library control) and the remainder was aliquoted and frozen for experiments. The M-ES Tn library was grown in MtbYM rich broth, then washed and diluted in PBS-T to OD_600_ = 0.05. Triplicate samples of the diluted M-ES Tn library were collected (Liquid control) and the diluted M-ES Tn library was plated on MtbYM rich agar in triplicate (Plate control) for Tn-seq. Mice were injected via the lateral tail vein with ∼10^5^ CFU of the M-ES Tn library to seed at least 10^4^ CFU in the lungs and spleen. Mice (*n*=3) were euthanized at days 1, 7, 21, and 42. Lung and spleen homogenates were plated on MtbYM rich agar to recover surviving Tn mutants for Tn-seq. **(B)** *M. tuberculosis* CFU recovered from lungs (blue) and spleens (green). **(C)** Heat map of log_2_ fold change in Tn mutant abundance between the indicated conditions determined by TnseqDiff analysis of Tn-seq data. Positive values (red) indicate a relative fitness advantage; negative values (blue) indicate a relative fitness defect. **(D-F)** Volcano plots of TnseqDiff analysis of Tn-seq data for the M-ES Tn mutant library for **(D)** lungs at day 1 compared to the Plate control; **(E)** lungs at day 21 compared to lungs at day 1; or **(F)** spleens at day 21 compared to spleens at day 1. Dashed lines indicate cutoffs for statistical significance of ± 2 log_2_ fold change and adjusted *P* value <0.025. Tn mutants meeting these significance cutoffs are colored and labeled.

To screen the M-ES Tn library for fitness defects in mice, the library was grown in MtbYM rich liquid, pelleted in triplicate for genomic DNA extraction as an input control (Liquid control), plated on MtbYM rich agar as a control for recovery on plates (Plate control), and used to infect C57BL/6J mice by intravenous injection (**Fig 2A**). The intravenous route was used to ensure adequate representation of all Tn mutants in both lung and spleen tissues. At days 1, 7, 21 and 42 post-infection, groups of mice (*n*=3) were euthanized and surviving *M. tuberculosis* CFU were recovered from lung and spleen homogenates by plating on MtbYM rich agar (**Fig 2A**). We noted high variability in *M. tuberculosis* CFU recovered from the lungs at days 7 and 42 (**Fig 2B**). For analysis of Tn mutant fitness by Tn-seq, we selected plates containing at least 10^4^ *M. tuberculosis* CFU for genomic DNA extraction. Since only one mouse each at the day 7 and day 42 time points had sufficient CFU in the lungs for Tn-seq analysis (**Fig 2B**), the lung samples were analyzed only at days 1 and 21. Sufficient bacteria were recovered from the spleens at all time points for Tn-seq analysis (**Fig 2B**).

To determine Tn mutant fitness, the triplicate genomic DNA samples from the library input, liquid input, plate input, and tissue homogenates plated on MtbYM rich agar (**Fig 2A**) were subjected to Tn-seq analysis and read counts for each Tn insertion site were tabulated (**S2 Table**). To determine relative Tn mutant fitness, we used TnseqDiff, which identifies conditionally essential genes between conditions based on the relative frequencies of Tn-seq sequencing reads at each Tn insertion site [41]. We previously used this approach to identify Tn mutants with altered fitness in low-complexity *M. tuberculosis* Tn mutant libraries treated with antibiotics *in vitro* [28]. Complete TnseqDiff analyses are provided in **S3 Table**.

No statistically significant fitness defects were observed in comparisons between the original M-ES Tn library and the MtbYM rich liquid culture that was used to infect the mice (**Fig 2C**, **S1A Fig**), suggesting that all Tn mutants grow equally well in MtbYM rich medium. In TnseqDiff comparisons between the original M-ES Tn library and the input plated on MtbYM rich agar, several M-ES Tn mutants showed slight, but statistically significant fitness defects (*purF*::Tn, *purM*::Tn, *purQ*::Tn, *gltB*::Tn, *panC*::Tn; **Fig 2C**, **S1B Fig**). These Tn mutants may have growth defects on MtbYM rich agar compared to other mutants in the M-ES Tn library, which causes their under-representation after recovery on plates. These data suggest that most M-ES Tn mutants are efficiently recovered on MtbYM rich agar and that any fitness defects observed following mouse infection are due to inability to survive within the mouse rather than a competitive disadvantage on MtbYM rich agar.

To determine relative Tn mutant fitness at day 1 post-infection, we compared Tn mutant abundance in plated mouse tissues to the input plated on MtbYM agar (D1 lungs/plate or D1 spleen/plate) using TnseqDiff. Three Tn mutants (*gltB*::Tn, *pheA*::Tn, *panC*::Tn) were significantly under-represented in the lungs at day 1 post-infection (**Figs 2C and 2D**). Attenuation of *panC*::Tn was expected, as *panC* was essential for *M. tuberculosis* growth in BALB/c mice [29]. We observed similarly reduced fitness of the *gltB*::Tn and *pheA*::Tn mutants in the spleen at day 1 post-infection (**Fig 2C**, **S1C Fig**). These data suggest that Phe biosynthesis (PheA) and Glu biosynthesis (GltB) are essential for *M. tuberculosis* survival within host tissues at the earliest stage of infection.

To determine Tn mutant fitness in the lungs and spleens at later times post-infection, the day 7, day 21 or day 42 outputs were compared to the day 1 input from the corresponding tissue using TnseqDiff. We identified 14 Tn mutants that exhibited significantly reduced fitness in the lungs at day 21 post-infection (**Figs 2C and 2E**). These included Tn mutants in genes encoding enzymes required for purine and thiamine biosynthesis (*purF*::Tn, *purM*::Tn, *purQ*::Tn), amino acid biosynthesis (*pheA*::Tn, *trpB*::Tn, *trpG*::Tn, *proC*::Tn), riboflavin synthesis (*ribA2*::Tn, *ribG*::Tn) and PABA synthesis (*pabB*::Tn), as well as the *panC*::Tn and *phoP*::Tn positive controls. Attenuation of the *trpB*::Tn and *trpG*::Tn mutants was expected as another gene in the Trp biosynthesis pathway (*trpE*) was previously implicated in *M. tuberculosis* growth in C57BL/6 mice [13].

We observed a similar pattern of attenuation in the spleens at day 21. Most Tn mutants that were attenuated in lungs also exhibited significant fitness defects in the spleen (**Figs 2C and 2F**). One exception is *pabB*::Tn, which was significantly attenuated in lungs but not spleens (**Figs 2C, 2E and 2F**). The Tn mutants that exhibited significant fitness defects in the spleen at day 21 were also significantly attenuated in the spleen at days 7 and 42 post-infection (**Fig 2C; S1D and S1E Figs**). The *pheA*::Tn mutant was significantly attenuated in lungs and spleens at all time points, despite significantly reduced fitness at day 1 (**Fig 2C**). In contrast, the *gltB*::Tn mutant was attenuated in lungs and spleens, but this did not achieve statistical significance, possibly because the *gltB*::Tn mutant was at much lower abundance in the day 1 normalization controls (**Fig 2C, S1C Fig**). Overall, these data suggest that *M. tuberculosis* requires multiple central metabolic pathways, particularly purine nucleotide and thiamine biosynthesis, riboflavin biosynthesis, and biosynthesis of specific amino acids (Phe, Trp, Pro, Glu) to replicate within host tissues.

While most M-ES Tn mutants with *in vitro* growth defects were attenuated in mice, the *ilvA*::Tn mutant was a notable exception. The *ilvA*::Tn mutant did not replicate in Middlebrook 7H9 *in vitro* (**Fig 1C**), but exhibited no statistically significant fitness defects in mouse lungs or spleens at any time point (**Fig 2C**). These data indicate that *M. tuberculosis* does not require the *de novo* Ile biosynthesis during mouse infection and suggest that it can acquire sufficient Ile from the host.

### Purine and thiamine auxotrophy is cidal for *M. tuberculosis in vitro* and in mice

To confirm the results of our Tn-seq screen, we selected two Tn mutants for individual retesting. We selected *purF*::Tn as a representative mutant with defects in biosynthesis of purine and thiamine. PurF catalyzes the first committed step of *de novo* purine nucleotide biosynthesis, which also produces a metabolic precursor required for the *de novo* synthesis of thiamine (vitamin B1) (**Fig 3A**) [42]. In *M. tuberculosis*, purine nucleotides can be produced either *de novo* or via a salvage pathway from hypoxanthine, which is converted to the purine nucleotide precursor inosine monophosphate (IMP) by hypoxanthine-guanine phosphoribosyltransferase (Hpt) [43].

**Fig 3.**
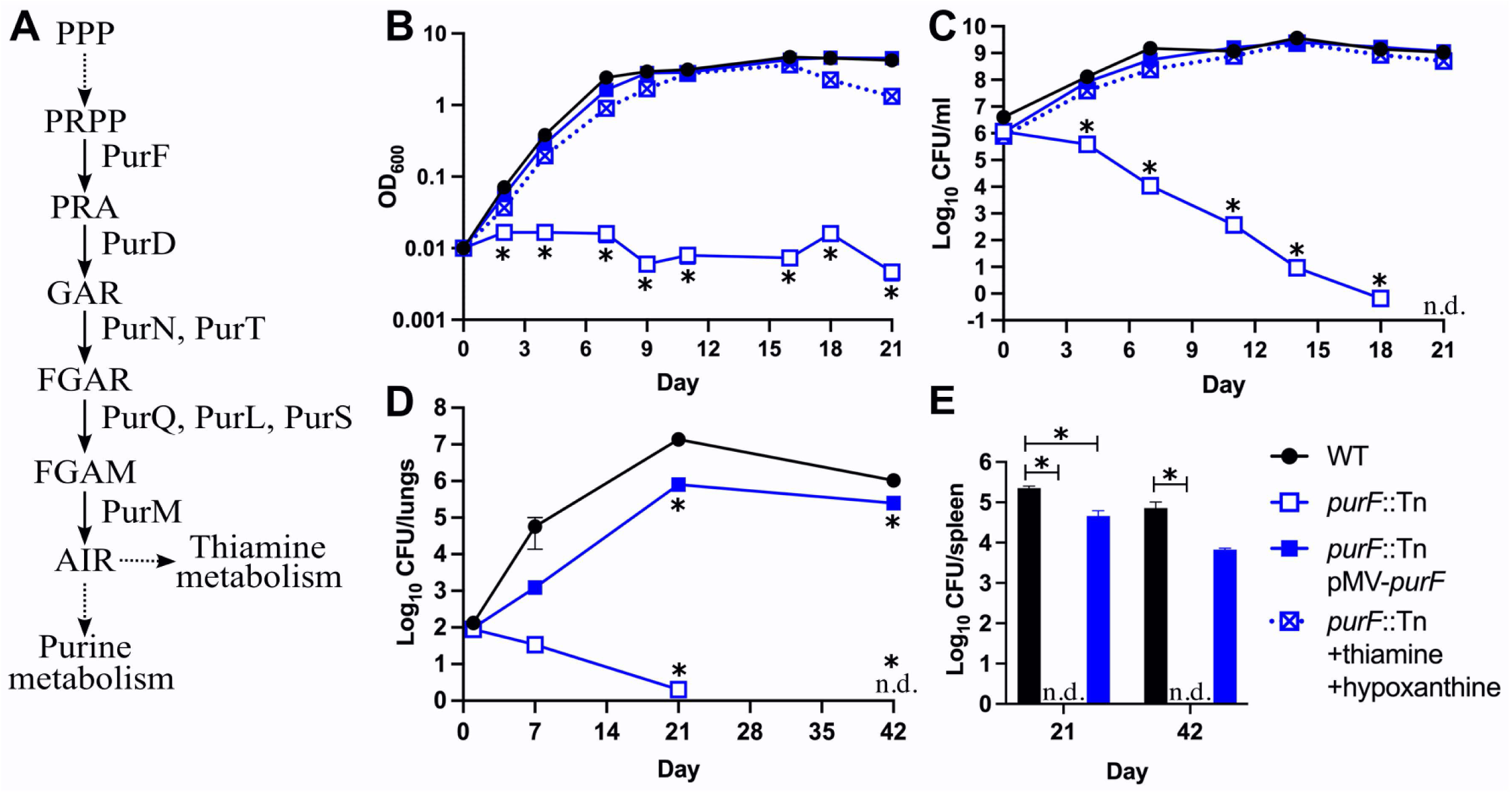
Loss of *purF* causes death of *M. tuberculosis* in the absence of nutrient supplementation *in vitro* and in mice. **(A)** Pathway for *de novo* synthesis of purine nucleotides and thiamine in *M. tuberculosis*. Solid lines indicate a single step; dashed lines indicate multiple steps. Proteins that catalyze the first five steps are indicated. Intermediate abbreviations: PPP, pentose phosphate pathway; PRPP, 5-phosphoribosyl pyrophosphate; PRA, 5-phospho-D-ribosylamine; GAR, 5’-phosphoribosylglycinamide; FGAR, 5’-phosphoribosyl-N-formylglycinamide; FGAM, 5’-phosphoribosyl N-formylglycinamidine; AIR, 5-aminoimidazole ribotide. **(B-C)** Strains grown in 7H9 supplemented with 60 μM thiamine and 150 μM hypoxanthine were washed in PBS-T and diluted to OD_600_ = 0.01 in 7H9 or 7H9 supplemented with 60 μM thiamine and 150 μM hypoxanthine. Bacterial growth and survival were measured by **(B)** optical density at 600 nm or **(C)** serial dilutions and plating to recover viable CFU. **(D-E)** C57BL/6J mice were infected by the aerosol with the indicated strains. Mice (*n*=6) were euthanized at days 1, 7, 21, and 42 post-infection. Lung **(D)** and spleen **(E)** homogenates were serially diluted and plated to recover viable CFU. In **(C-E)**, WT and *purF*::Tn pMV-*purF* were plated on 7H10 agar; *purF*::Tn was plated on 7H10 agar supplemented with 60 μM thiamine and 150 μM hypoxanthine. Data represent the mean ± standard error of three biological replicates **(B-C)** or six animals **(D-E)**. Asterisks indicate *P*-value < 0.05; n.d. indicates not detected (detection limit = 1 CFU).

We confirmed that the *purF*::Tn mutant fails to grow in standard Middlebrook 7H9 medium and that the growth defect can be alleviated with exogenous hypoxanthine and thiamine (**Fig 3B**), the two components of MtbYM rich predicted to bypass *de novo* purine biosynthesis [27]. Exogenous hypoxanthine alone partially rescued growth of the *purF*::Tn mutant, but full growth restoration only occurred with both nutrients added (**S2 Fig**). The growth defect of the *purF*::Tn mutant in 7H9 was also fully complemented by *purF* expressed from a plasmid (pMV-*purF*, **Fig 3B**). The *purF*::Tn mutant lost viability in unsupplemented 7H9, with viable CFU below the limit of detection (1 CFU/ml) after 21 days of incubation (**Fig 3C**). Viability of the *purF*::Tn mutant was restored by either exogenous hypoxanthine and thiamine or by complementation with pMV-*purF* (**Fig 3C**). These data demonstrate that loss of PurF function causes death of *M. tuberculosis* in the absence of nutrient supplementation.

To determine if *M. tuberculosis* requires PurF during infection, we infected C57BL/6J mice by aerosol with WT Erdman, *purF*::Tn or the *purF*::Tn pMV-*purF* complemented strain. All strains were confirmed to produce the PDIM lipid that is essential for virulence using thin layer chromatography of ^14^C-propionate labelled lipid extracts (**S3 Fig**). The *purF*::Tn mutant established infection in the lungs at day 1 post-infection, but it failed to replicate, was cleared from the lungs by day 42 post-infection, and failed to disseminate to the spleen (**Figs 3D and 3E**). Attenuation of the *purF*::Tn mutant was partially complemented by pMV-*purF*. The *purF*::Tn pMV-*purF* strain replicated in the lungs and disseminated to spleen, but significantly fewer CFU were recovered compared to the WT control (**Figs 3D and 3E**). Expression of *purF* from the pMV-*purF* plasmid may be insufficient to support *M. tuberculosis* growth in host tissues. Alternatively, the *purF*::Tn insertion may be polar on the 3’ *purM* gene, which is also a M-ES gene [11]. Reduced expression of *purF* or *purM* could prevent normal *M. tuberculosis* replication in mice, in which demand for purine nucleotides and/or thiamine may be higher as compared to *in vitro* culture. Overall, our data demonstrate that the Tn insertion in *purF*, not a secondary mutation, causes attenuation in mice. These data indicate that *M. tuberculosis* cannot obtain sufficient purine nucleotide precursors and/or thiamine to support growth in host tissues and implicate the purine and thiamine biosynthetic pathways as TB drug targets.

### *M. tuberculosis* requires Phe biosynthesis for survival *in vitro* and in mice

We selected the *pheA*::Tn mutant for retesting as the role of Phe biosynthesis in *M. tuberculosis* fitness either *in vitro* or in the host has not previously been tested and the Phe biosynthesis pathway is absent in humans, making it an attractive drug target [44]. PheA is a prephenate dehydratase required for both pathways of Phe synthesis (**Fig 4A**). PheA catalyzes the first step in Phe synthesis from prephenate [45], which is produced from chorismate by chorismate mutase [44]. Two *M. tuberculosis* proteins have chorismate mutase activity (Rv1885c and Rv0948c), but Rv1885c is secreted while Rv0948c is cytosolic [46-48]. Only *rv0948c* is a M-ES gene [11], suggesting that it functions as the chorismate mutase in Phe synthesis. In plants and some bacteria, Phe synthesis can also proceed via an L-arogenate intermediate [44]. L-arogenate has been detected in *M. tuberculosis* [49], and the aminotransferase Rv1178c may synthesize L-arogenate from chorismate, as this enzyme activity has been demonstrated *in vitro* [50]. PheA is predicted to act as the dehydratase to convert L-arogenate to Phe.

**Fig 4.**
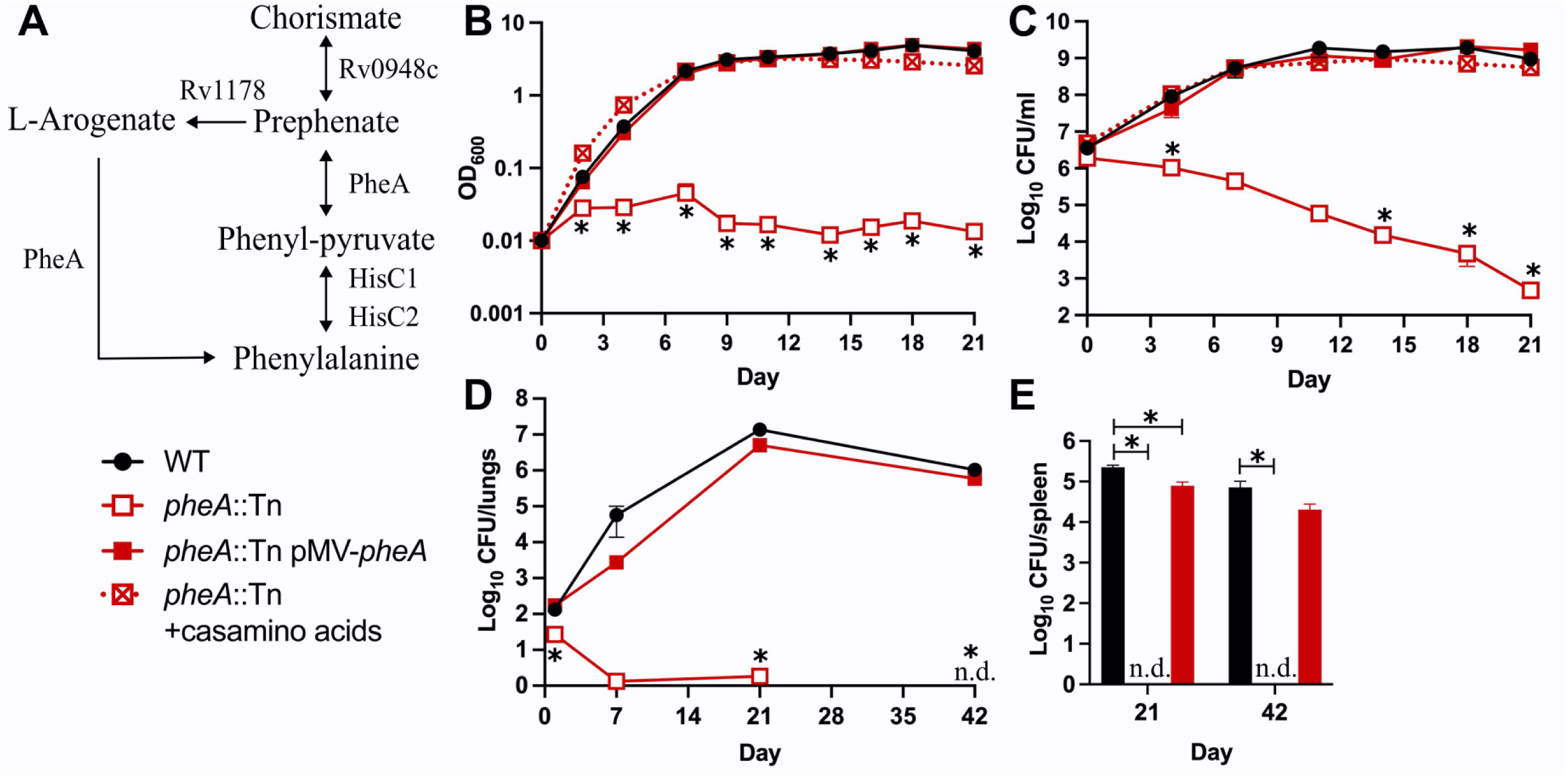
*M. tuberculosis* requires Phe synthesis for survival *in vitro* in the absence of amino acid supplementation and in mice. **(A)** Phenylalanine biosynthesis in *M. tuberculosis*. Single arrows represent one-way reactions. Double arrows represent reversible reactions. Proteins predicted to catalyze each step are indicated. **(B-C)** Strains grown in 7H9 supplemented with 0.5% casamino acids were washed in PBS-T and diluted to OD_600_ = 0.01 in 7H9 or 7H9 supplemented with 0.5% casamino acids. Bacterial growth and survival were measured by **(B)** optical density at 600 nm and **(C)** serial dilutions and plating to recover viable CFU. **(D-E)** C57BL/6J mice were infected by aerosol with the indicated strains. Mice (*n*=6) were euthanized at days 1, 7, 21, and 42 post-infection. Lung **(D)** and spleen **(E)** homogenates were serially diluted and plated to recover viable CFU. In **(C-E)**, WT and *pheA*::Tn pMV-*pheA* were plated on 7H10 agar; *pheA*::Tn was plated on 7H10 agar with 0.5% casamino acids. Data represent the mean ± standard error of three biological replicates **(B-C)** or six animals **(D-E)**. Asterisks indicate *P*-value < 0.05; n.d. indicates not detected (detection limit = 1 CFU).

We confirmed that the *pheA*::Tn mutant does not grow in 7H9 medium unless the medium is supplemented with casamino acids, which contains Phe (**Fig 4B**). The growth defect of the *pheA*::Tn mutant in 7H9 was also fully complemented with plasmid-encoded *pheA* (pMV-*pheA*, **Fig 4B**). The *pheA*::Tn mutant also lost viability in 7H9 in the absence of casamino acid supplementation, indicating that an inability to acquire or produce Phe is cidal to *M. tuberculosis* (**Fig 4C**). This survival defect was also fully complemented by pMV-*pheA* (**Fig 4C**). These data demonstrate that blocking Phe synthesis causes death of *M. tuberculosis* in the absence of exogenous nutrient supplementation.

To confirm attenuation of the *pheA*::Tn mutant, C57BL/6J mice were infected by aerosol with WT Erdman, *pheA*::Tn, or the *pheA*::Tn pMV-*pheA* complemented strain. The *pheA*::Tn and *pheA*::Tn pMV-*pheA* strains both produce the PDIM lipid (**Fig S3**). Despite using a higher inoculating dose, we observed significantly fewer *pheA*::Tn bacteria in the lungs at 24 hr post-infection compared to either the WT or complemented controls (**Fig 4D**). This is consistent with the significant reduction in fitness we observed for the *pheA*::Tn mutant at day 1 post-infection compared to the plated input control in our Tn-seq screen (**Figs 2C & 2D**). These data suggest that PheA is required for *M. tuberculosis* to establish infection in the lungs. The *pheA*::Tn mutant was also incapable of replicating in mouse lungs, was cleared to below the limit of detection from lung tissues by 42 days post-infection, and failed to disseminate to the spleen (**Figs 4D & 4E**). The *pheA*::Tn pMV-*pheA* complemented strain replicated in the lungs (**Fig 4D**) and disseminated to spleen (**Fig 4E**) similarly to the WT control. These data show that attenuation of the *pheA*::Tn mutant is due to the Tn insertion in *pheA* and not a secondary mutation. These data also demonstrate that *M. tuberculosis* requires Phe biosynthesis to survive within the host, suggest that Phe is not available in sufficient quantities in host tissues to support *M. tuberculosis* growth, and indicate that PheA is a potential target for development of new TB drugs.

## DISCUSSION

Most metabolic pathways that are essential for *M. tuberculosis* growth in standard *in vitro* culture conditions have not previously been evaluated to determine their importance during infection because mutant strains lacking these metabolic functions cannot be generated using standard methods. Here we use a Tn mutant collection generated using a nutrient-enriched growth medium to demonstrate that several metabolic pathways essential for growth in standard *in vitro* culture conditions are also essential in the host using a mouse infection model. Our data show that *M. tuberculosis* requires *de novo* purine and thiamine biosynthesis as well as Phe biosynthesis for replication in the lungs and dissemination to the spleen in aerosol-infected mice. In addition, we show that an inability to synthesize Phe or purine and thiamine causes death of *M. tuberculosis* in the absence of exogenous nutrients, suggesting that inhibitors of these biosynthetic pathways would be bacteriocidal. Therefore, our data implicate the Phe and purine/thiamine biosynthesis pathways as promising targets for development of novel anti-tubercular drugs.

Phe biosynthesis has been explored as a target for anti-tubercular drug development because mammals lack *de novo* Phe synthesis, which is expected to limit the toxicity of compounds targeting this pathway [44]. Our data support PheA as a potential drug target since our *pheA*::Tn mutant exhibited strong attenuation in the mouse model, suggesting that *M. tuberculosis* cannot acquire sufficient Phe from the host. Phe biosynthesis also requires conversion of chorismate to prephenate by chorismate mutase [44]. Inhibitors of *M. tuberculosis* chorismate mutase have been identified, but most of these compounds had high minimal inhibitory concentrations (MICs), suggesting that they have low whole cell permeability [51-53]. Our data suggest that *M. tuberculosis* can import Phe, since growth of the *pheA*::Tn mutant *in vitro* was restored by amino acid supplementation. Therefore, it may be possible to design PheA or chorismate mutase inhibitors with improved permeability properties for use as anti-tubercular agents.

Purine nucleotide biosynthesis is critical for the virulence of many pathogens and has also been explored as a drug development target [54]. Although mammals also synthesize purine nucleotides *de novo*, the enzymes are structurally distinct, suggesting that specific inhibitors of bacterial enzymes can be designed [42]. Our data demonstrate that *M. tuberculosis* requires PurF, which catalyzes the first committed step of *de novo* purine nucleotide and thiamine co-factor biosynthesis, for growth in the host. Our data suggest that both purine and thiamine biosynthesis are critical functions of PurF, as *in vitro* growth was fully restored only by adding exogenous thiamine and hypoxanthine. Attenuation of the *purF*::Tn mutant in mice was only partially reversed by complementation, despite full restoration of *in vitro* growth in the absence of nutrient supplementation. This suggests that the demand for purine and/or thiamine biosynthesis is greater within host tissues and that even partial loss of PurF function can limit *M. tuberculosis* growth. Collectively, our data suggest that *M. tuberculosis* cannot acquire sufficient purine nucleotides and/or thiamine from host tissues to support its growth.

Our data also indicate that *M. tuberculosis* requires other enzymes in the purine and thiamine biosynthetic pathway for growth in the host, as *purM*::Tn and *purQ*::Tn mutants were highly attenuated in our Tn-seq screen. Our data are consistent with a prior report that a *M. tuberculosis purC* mutant was cleared from the tissues of intravenously-infected mice [55]. Our data are also consistent with the observation that depletion of GuaB2, which is required for *de novo* synthesis of guanine nucleotides, prevents *M. tuberculosis* replication in aerosol-infected mice [56]. Loss of either PurC or GuaB2 function is predicted to prevent production of guanine nucleotides, but not thiamine. Whether an inability to produce thiamine contributes to the attenuation of the *purF*::Tn mutant remains to be determined. In addition, it is unclear whether *M. tuberculosis* requires *de novo* purine and/or thiamine synthesis only for replication during acute infection or also for persistence in host tissues during chronic infection. Our future studies will explore these questions using *M. tuberculosis* strains that conditionally express enzymes specific to the thiamine or purine biosynthetic pathways or that are required for both pathways, like PurF. Overall, our data are consistent with prior studies indicating that *M. tuberculosis* requires *de novo* purine biosynthesis for growth in the host and support further development of inhibitors targeting this pathway.

Nucleotide synthesis inhibitors, which include approved cancer chemotherapeutics, have shown efficacy against other bacterial pathogens, but can be toxic to mammalian cells [54]. Structures of *M. tuberculosis* PurN, PurH, and PurF, which are involved in *de novo* purine synthesis, revealed structural distinctions from the equivalent human enzymes, suggesting that specific inhibitors could be designed against these proteins [57-59]. *Mycobacterium abscessus* PurC is also structurally distinct from the human ortholog, and PurC inhibitors that prevent *M. tuberculosis* growth *in vitro* were recently described [60]. An indazole sulfonamide compound that inhibits GuaB2 was cidal against replicating *M. tuberculosis in vitro* but lacked activity in infected mice due to either low drug concentration in lung lesions, slower growth of *M. tuberculosis* in chronically infected mice, or high levels of guanine in host tissues that antagonize the drug [61]. It may be possible to develop improved drugs targeting other enzymes in the purine biosynthetic pathway that would disrupt both purine and thiamine biosynthesis. As *de novo* purine biosynthesis includes 10 enzymatic steps, it represents a pathway rich in targets for the development of novel anti-tubercular agents.

The *de novo* thiamine biosynthesis pathway is also a promising target for development of novel antimicrobials given that it is absent in humans, who must obtain thiamine in the diet [62]. Inhibitors of thiamine biosynthesis with *in vitro* efficacy have been reported for *Salmonella* and *Pseudomonas* [63, 64]. Inhibitors of *M. tuberculosis* ThiE, which catalyzes the final step in production of the active co-enzyme thiamine pyrophosphate, have also been identified, one of which potently inhibited *in vitro* growth [65]. However, this study did not test whether the compound specifically targeted ThiE through either selection of resistant mutants or use of hyper-morphic strains with reduced ThiE expression. Further studies will be needed to determine if *M. tuberculosis* requires thiamine synthesis during infection and to identify the steps in the biosynthetic pathway that are most vulnerable to inhibition.

Loss of either PheA or PurF function causes rapid death of *M. tuberculosis* in the absence of exogenous nutrient supplementation, suggesting that inhibitors targeting these enzymes would be bacteriocidal. The *pheA*::Tn mutant died *in vitro* at a rate comparable to a methionine auxotroph (Δ*metA*) without amino acid supplementation [18]. The *purF*::Tn mutant was sterilized to below the limit of detection after only 21 days of culture without hypoxanthine and thiamine. This is a faster rate of death in the absence of nutrient supplementation than any other auxotrophic *M. tuberculosis* strains that have been characterized, including Arg biosynthesis mutants, which are sterilized by Arg deprivation [18, 19]. However, the molecular mechanisms that cause death of either the *pheA*::Tn or *purF*::Tn mutants remain to be determined.

For other auxotrophic *M. tuberculosis* mutants, loss of viability was correlated with multiple effects on microbial physiology. For example, the Δ*metA* mutant cannot produce either Met or *S*-adenosylmethionine (SAM), an essential co-factor in one-carbon metabolism [18]. Similarly, death of Arg biosynthesis mutants deprived of Arg was correlated with increased production of endogenous reactive oxygen species and DNA damage, in addition to reduced intracellular Arg [19]. Reduced Phe production in the *pheA*::Tn mutant may deplete chorismate because chorismate mutase is feedback inhibited by Phe [66]. Chorismate is also required as a precursor for synthesis of Trp, menaquinones and the mycobactin siderophore, so its depletion would be expected to affect multiple processes [47]. Loss of PurF function may be cidal in the absence of nutrient supplementation due to reduced production of both purine nucleotides, which are needed for DNA synthesis, RNA synthesis and energy storage [54], and thiamine, which is needed for activity of multiple enzymes involved in central carbon metabolism and branched chain amino acid biosynthesis [62]. Alternatively, the *purF*::Tn mutant may be hypersusceptible to exogenous stress, including oxygen limitation, as previously reported for a *Mycobacterium smegmatis purF* mutant [67]. The mechanisms by which these mutations cause death of *M. tuberculosis* will be explored in our future studies.

In addition to the *purF*::Tn and *pheA*::Tn mutants that we selected for follow-up studies, we identified several other Tn mutants that exhibited strong fitness defects in mice. These included Tn insertions in genes required for riboflavin synthesis (*ribA2*, *ribG*), PABA synthesis (*pabB*), and synthesis of the amino acids Pro and Glu (*proC, gltB*). Our data suggest that *M. tuberculosis* cannot acquire these nutrients in sufficient quantities from host tissues and must synthesize them *de novo*. To prioritize these conditionally essential metabolic pathways as targets for drug development, we will consider whether the pathway is present in mammalian cells, whether loss of the pathway is cidal *in vitro* in the absence of nutrient supplementation, and whether the pathway is vulnerable to inhibition.

Riboflavin, Glu and PABA synthesis pathways are absent from mammalian cells, so compounds targeting these pathways would be expected to have low toxicity. We can use our Tn mutants to assess whether starvation for these nutrients is cidal for *M. tuberculosis in vitro*, as we have done for the *pheA*::Tn and *purF*::Tn mutants. PABA is a metabolic precursor in folate biosynthesis. A *M. tuberculosis* PabB inhibitor has been reported to overcome resistance to anti-folate drugs [68], indicating that compounds targeting PabB can be developed. Vulnerability of *M. tuberculosis* to loss of gene function *in vitro* has been estimated based upon reduced fitness with transcriptional inhibition using CRISPR interference (CRISPRi) [9]. A genome-wide CRISPRi screen revealed that *ribA2* and *gltB* were among the genes most vulnerable to inhibition, with vulnerability indices in the top 4% of all of *M. tuberculosis* genes [9]. Many other M-ES genes that we analyzed including *pheA*, *purF* and other genes required for purine biosynthesis were also highly vulnerable, with vulnerability indices similar to the targets of existing anti-TB drugs [9]. Our data indicate that riboflavin, Glu and PABA synthesis are all required for acute *M. tuberculosis* infection and should be considered as potential targets for development of new anti-tubercular agents. Future studies will address whether these genes are also essential during chronic *M. tuberculosis* infection using conditional expression strains.

While our data indicate that *M. tuberculosis* requires most of the M-ES genes that we analyzed for survival in mice, *ilvA* was an exception. IlvA catalyzes the first step in *de novo* synthesis of isoleucine (Ile), a branched chain amino acid (BCAA) [69]. BCAA biosynthesis has been actively investigated as a target for antimicrobial development because these pathways are absent in mammals [70]. In addition, *M. tuberculosis ilvA* is highly vulnerable to inhibition *in vitro* [9]. However, our data suggest that IlvA would be a poor drug target because the *ilvA*::Tn mutant was not attenuated in mice, despite strong *in vitro* growth inhibition. Our data suggest that *M. tuberculosis* can obtain sufficient Ile from host tissues to support its growth. Our data are consistent with the observation that a *M. tuberculosis* mutant lacking *ilvB1*, which is required for synthesis of all BCAAs (Ile, Leu, Val), was not cleared from lung or spleen tissues in intravenously-infected mice, despite loss of viability *in vitro* without exogenous BCAA [71]. Indeed *M. tuberculosis* can acquire all BCAAs directly from infected macrophages [72]. Our data highlight the importance of directly testing whether the biosynthetic pathways essential for *in vitro* growth are also necessary during infection prior to pursuing these pathways as drug development targets.

Finally, our work highlights a new strategy for identifying *M. tuberculosis* mutants that are attenuated during mouse infection using defined pools of Tn mutants and quantification of Tn mutant fitness with deep sequencing. Most genome-wide screens of *M. tuberculosis* Tn mutants in the mouse model used complex Tn mutant pools, which enabled analysis of Tn mutant fitness only in mouse spleens due to a strict colonization bottleneck in the lungs [12-14, 16]. Initial Tn mutant screens conducted with signature-tagged mutagenesis technology used low-complexity Tn mutant pools to identify *M. tuberculosis* Tn mutants specifically attenuated in the lungs [17, 73, 74]. A few recent studies have examined fitness of deletion mutants with sequence barcodes and quantification of mutant fitness in the lungs by sequencing [75, 76]. However, to our knowledge this is the first study to use low-complexity Tn mutant pools and Tn-seq to identify *M. tuberculosis* mutants attenuated in mouse lungs. A similar strategy combined with high-dose aerosol infection could be used to identify *M. tuberculosis* genes that are required for aerosol transmission and for fitness within lung tissues.

Our work is also the first to determine role of multiple essential central metabolic pathways in *M. tuberculosis* pathogenesis. Our results directly demonstrate that *M. tuberculosis* requires both *de novo* Phe synthesis and *de novo* purine/thiamine synthesis for replication and survival in infected mice, implicating these pathways as prime targets for development of new TB drugs. Our Tn-seq screen identified IlvA as a conditionally essential gene that is not required for *M. tuberculosis* survival in the host, suggesting that BCAA biosynthesis pathways should be de-prioritized for drug development. Our screen also revealed several other metabolic pathways, including synthesis of other amino acids (Glu, Pro), PABA and riboflavin that *M. tuberculosis* requires for growth in host tissues. Overall, our results widen the scope of potential targets for TB drug development.

## MATERIALS AND METHODS

### Bacterial strains and growth conditions

*M. tuberculosis* Erdman wild-type and derivative strains were grown aerobically at 37°C in Middlebrook 7H9 (Difco) liquid medium supplemented with 10% albumin-dextrose-saline (ADS), 0.5% glycerol, and 0.1% Tween-80 or on Middlebrook 7H10 (Difco) agar supplemented with 10% oleic acid-albumin-dextrose-catalase (OADC; BD Biosciences) and 0.5% glycerol unless otherwise noted. Frozen stocks of *M. tuberculosis* strains were made by growing cultures to late-exponential phase, adding glycerol to 15% final concentration, and storing at -80°C. Liquid cultures of M-ES Tn mutants or the M-ES Tn library were grown in MtbYM rich medium (MtbYM) pH 6.6 [27] supplemented with 10% OADC and 0.05% tyloxapol or on MtbYM agar plates [27], unless otherwise noted. Antimicrobials were used at the following concentrations: kanamycin (Kan) 25 μg/ml for agar or 15 μg/ml for liquid, hygromycin (Hyg) 50 μg/ml, cycloheximide 100 μg/ml.

### Recovery of Tn mutants from the arrayed Tn mutant library

Tn mutants were recovered from the mapped location in our arrayed Tn mutant library [28] by streaking on MtbYM agar containing Kan and incubating at 37˚C for at least three weeks. Up to four individual colonies were picked and grown in 10 ml of MtbYM broth containing Kan at 37˚C with aeration. The Tn insertion site was confirmed by PCR using a gene-specific primer 5’ or 3’ of the TA site and a primer specific to the Tn (**Table S4**) followed by Sanger sequencing of the PCR product. M-ES Tn mutants were screened for secondary point mutations that disrupt PDIM production known to be present in our Tn library by PCR and Sanger sequencing. We screened for mutations in *ppsD* and *ppsE* previously identified in other Tn mutants [28] and for additional common mutations in *ppsB*, *ppsC* and *ppsE* identified by whole-genome sequencing of M-ES Tn mutants (**Table S4**).

### Whole genome sequencing

The whole genome of each M-ES Tn mutant included in the M-ES library was sequenced by short-read IIlumina sequencing to identify mutations in the phthiocerol dimycocerosate (PDIM) locus that could result in PDIM deficiency. Tn mutants were grown to an OD_600_ of 1.0 in MtbYM broth; genomic DNA (gDNA) was extracted by the CTAB-lysozyme method [77] and cleaned with the Genomic DNA concentrator and cleanup kit 25 (Zymo). gDNA was submitted to SeqCenter (formerly Microbial Genome Sequencing Center, MiGS) (Pittsburgh, PA) for library preparation and Illumina sequencing (151 bp paired-end output, 400 Mbp, 2.67 M reads per sample). To generate a consensus sequence for each strain, paired reads were mapped to the *M. tuberculosis* Erdman reference genome (NC_020559.1) using the “map to reference” function in Geneious 2021 software (Biomatters, Ltd.) as previously described [28]. To identify single nucleotide polymorphisms (SNPs) in Tn mutant genomes, the WT Erdman and Tn mutant consensus sequences were aligned in Geneious using the “Align Whole Genomes” function with the default Mauve Genome parameters, as described [28]. Tn mutant genomes were only examined for SNPs in the PDIM locus (*tesA – lppX*). SNPs identified in genes required for PDIM synthesis were confirmed by PCR amplification and Sanger sequencing (**Table S4**). Tn mutants with confirmed SNPs in the PDIM locus were excluded from the study. To confirm Tn insertion sites in the Tn mutants, reads were mapped to the *Himar1* Tn sequence in Geneious as described above and sequences adjacent to the Tn were compared to the *M. tuberculosis* Erdman reference genome. Mutants with more than one Tn insertion were excluded from the study.

### Creation and screening of the M-ES Tn library by Tn-seq

We created our M-ES Tn mutant library using Tn mutants in genes predicted to be conditionally essential in Middlebrook 7H9 medium but not MtbYM rich medium (M-ES genes) [11, 27], with a single Tn insertion and no mutations in the PDIM locus based on whole-genome sequencing (**Table S1**). For positive controls, we isolated Tn mutants in genes known to be required for *M. tuberculosis* virulence (*panC*::Tn and *phoP*::Tn) [29, 36, 37]. For negative controls, we isolated Tn mutants in genes known to be dispensable during mouse infection (*rpfA*::Tn, *rpfE*::Tn, *tadA*::Tn, *pstB*::Tn, *pstC1*::Tn) [38-40]. Each Tn mutant was grown individually in MtbYM broth until late-logarithmic phase (OD_600_ ∼1.0) and mixed in equal abundance to make the M-ES Tn library. Frozen stocks of the M-ES Tn library were made by adding glycerol to 15% final concentration and freezing 1 ml aliquots. The remaining Tn library culture was aliquoted into three 10 ml cultures and pelleted by centrifugation (4100 x*g*, 10 min) for gDNA extraction as a total library (Library) control. For the Tn-seq screen, the Tn library was grown from a frozen stock to mid-exponential phase (OD_600_ = 0.5) in MtbYM broth, washed once with PBS containing 0.05% Tween-80 (PBS-T), and resuspended at OD_600_ = 0.05 in PBS-T. Three 9 ml samples of the diluted M-ES Tn library (∼10^7^ CFU) were pelleted by centrifugation (4100 x*g*, 10 min) for gDNA extraction as an input (Liquid) control. The diluted M-ES Tn library was serially diluted and plated on MtbYM agar to enumerate total bacterial CFU and to isolate a plated input (Plate) control.

C57BL/6J mice 7 weeks of age were purchased from Jackson Laboratory, USA. Mice were injected with 200 μl of the diluted M-ES Tn library (∼2 x10^5^ CFU) via the lateral tail vein to deliver ∼10,000 CFU to the mouse lungs. Groups of mice (*n*=3) were euthanized by CO_2_ overdose at days 1, 7, 21, and 42 post-infection. Lungs and spleens were collected, homogenized in PBS-T, serially diluted, and plated on MtbYM agar containing Kan and cycloheximide to enumerate total bacterial CFU in the sample and to recover at least 10^4^ CFU per plate for Tn-seq analysis. Plates were incubated at 37˚C with 5% CO_2_ until the biomass on the agar was confluent, up to four weeks. Plates with at least 10^4^ CFU for the Plate controls and mouse tissue homogenates were flooded with 2 ml of GTE buffer [77], and gently scraped with a plastic 10 μl loop to loosen the biomass. Bacteria were collected by centrifugation (4100 x*g*, 10 min). Genomic DNA was extracted from all input controls (Library, Liquid, Plate) and from plated mouse lung and spleen homogenate samples by the CTAB-lysozyme method [77], cleaned with the Genomic DNA clean and concentrator kit (Zymo), and submitted to the University of Minnesota Genomics Center (UMGC) for Tn-seq library preparation and Illumina sequencing.

### Transposon sequencing (Tn-seq) and data analysis

Tn-seq was performed as previously described [27]. *M. tuberculosis* gDNA was fragmented with a Covaris S220 ultrasonicator and a whole genome library was prepared using the TruSeq Nano library preparation kit (Illumina). Library fragments containing Tn junctions were PCR-amplified from the whole genome library using the Tn-specific primer Mariner_1R_TnSeq_noMm and Illumina p7 primer (**Table S4**). The amplified products were uniquely indexed to allow sample pooling and multiplexed sequencing. A total of 27 Mariner-enriched Tn-seq libraries were created. All libraries were pooled and sequenced on a NextSeq 2K P1 2×150-bp run (Illumina) with 20% PhiX target spiked in to account for the low diversity of Tn-seq libraries. Approximately 117 M pass filter reads were generated for the lane. All expected barcodes were detected and reads were balanced (Mean reads ≈ 2.5 M) with mean quality scores for all libraries ≥Q30.

Tn-seq analysis was done as previously described [28]. Sequencing reads were filtered to remove reads without the Tn sequence “GGACTTATCAGCCAACCTGT”. The 5’ Illumina adaptor sequences were trimmed using BBDuk (https://sourceforge.net/projects/bbmap/). Each trimmed read was cut to 30 bases and sequences not starting with TA were removed. Remaining reads were mapped to the *M. tuberculosis* Erdman genome (NC_020559.1) using HISAT2. Mapped reads were counted at each TA insertion site to generate read count tables for TnseqDiff analysis. TnseqDiff normalized the read counts using the default trimmed mean of M values (TMM) normalization method [78, 79], and then determined conditional essentiality for each TA insertion site between experimental conditions. TnseqDiff calculated the fold change and corresponding two-sided *P*-value for each TA insertion site [41]. All *P*-values were adjusted for multiple testing using the Benjamini-Hochberg procedure in TnseqDiff. The cut-off values for statistical significance were set at a fold-change of > log_2_ ±2 and an adjusted *P*-value < 0.025.

### Tn mutant complementation

Complementation vectors pMV-*purF* and pMV-*pheA* were in made in the integrating plasmid pMV306hyg [28] by Gibson assembly. Each gene was PCR-amplified (Phusion; New England Biolabs) along with ∼300 bases 5’ of the translation start site to include the native promoter using primers designed with 16- 20 bases complementary to the gene of interest and 18-22 bases that overlap the NcoI site in pMV306hyg (**Table S4**). PCR products were purified (Qiagen PCR purification kit) and assembled with pMV306hyg that was linearized by digestion with NcoI using HiFi DNA Assembly Master Mix (New England Biolabs) according to the manufacturer’s instructions followed by Sanger sequencing of the cloned gene. The *purF*::Tn and *pheA*::Tn mutants were electroporated with the corresponding complementation vector as described [80]. Each Tn mutant was grown in 7H9 with the appropriate supplements added (60 μM thiamine and 150 μM hypoxanthine for *purF*::Tn or 0.5% casamino acids for *pheA*::Tn). Transformants were selected on Middlebrook 7H10 agar containing Kan and Hyg without supplements. The complementing plasmid was confirmed by PCR (**Table S4**).

### Growth curves

Each M-ES Tn mutant in the M-ES Tn library and WT Erdman were grown individually in MtbYM broth to mid-exponential phase (OD_600_ = 0.4-0.6), pelleted by centrifugation (2850 x*g*, 10 min) and washed three times with PBS-T before diluting to OD_600_ = 0.01 in 7H9. Optical density (OD_600_) was measured every 2-3 days.

To confirm conditional essentiality of *purF*::Tn in 7H9, WT, *purF*::Tn and *purF*::Tn pMV-*purF* were grown to mid-exponential phase (OD_600_ = 0.4-0.6) in 7H9 supplemented with 60 μM thiamine and 150 μM hypoxanthine (7H9 +thiam +hypo), pelleted by centrifugation (2850 x*g*, 10 min) and washed three times with PBS-T before diluting to OD_600_ = 0.01 in 7H9 or 7H9 +thiam +hypo. The OD_600_ was measured every 2-3 days. Surviving bacteria were enumerated by serially diluting in PBS-T and plating on Middlebrook 7H10 agar with 10% OADC and 0.5% glycerol. 7H10 agar was additionally supplemented with 60 μM thiamine and 150 μM hypoxanthine for recovery of the *purF*::Tn mutant. At later time points, bacteria in 1 ml of the *purF*::Tn 7H9 culture were collected by centrifugation (3000 x*g*, 10 min), resuspended in 100 μl of PBS-T and plated. Plates were incubated for at least 4 weeks at 37˚C before enumerating CFU.

To confirm conditional essentiality of *pheA*::Tn in 7H9, WT, *pheA*::Tn and *pheA*::Tn pMV-*pheA* were grown to mid-exponential phase (OD_600_ = 0.4-0.6) in 7H9 supplemented with 0.5% casamino acids (7H9 +cas), pelleted by centrifugation (2850 x*g*, 10 min) and washed three times with PBS-T before diluting to OD_600_ = 0.01 in 7H9 or 7H9 +cas. The OD_600_ was measured every 2-3 days. Surviving bacteria were enumerated by serially diluting in PBS-T and plating on Middlebrook 7H10 agar with 10% OADC and 0.5% glycerol. 7H10 agar was additionally supplemented with 0.5% casamino acids for recovery of the *pheA*::Tn mutant. Plates were incubated for at least 4 weeks at 37˚C before enumerating CFU.

### Phthiocerol dimycocerosate (PDIM) labeling and detection

PDIM production was measured by a radiolabeling and thin layer chromatography (TLC) method [81]. Cultures grown to mid-logarithmic phase in 10 ml of 7H9 broth with appropriate supplements (7H9 +thiam +hypo for *purF*::Tn, 7H9 +cas for *pheA*::Tn) were labeled for 48 hr with 10 μCi of [1-^14^C] propionic acid, sodium salt (American Radiolabeled Chemicals, Inc; specific activity 50-60 mCi/mmol) prior to apolar lipid extraction and detection of the PDIM lipid by TLC as described [28].

### Mouse aerosol infection

For retesting individual Tn mutant and complemented strains, male and female C57BL/6J mice 6-8 weeks of age were purchased from Jackson Laboratory, USA, and infected via the aerosol route using an Inhalation Exposure System (GlasCol) as described [82]. The nebulizer was loaded with a bacterial suspension in PBS-T at OD_600_ = 0.007 for WT and complemented strains or OD_600_ = 0.01 for the *purF*::Tn or *pheA*::Tn mutant strains to deliver ∼100 CFU to the lungs. Groups of mice (*n* = 6, 3 female and 3 male) were euthanized by CO_2_ overdose at 1, 7, 21, and 42 days post-infection. Bacterial CFU were enumerated by plating serially diluted lung and spleen homogenates on 7H10 agar with cycloheximide. The 7H10 agar was supplemented with 60 μM thiamine and 150 μM hypoxanthine for recovery of the *purF*::Tn mutant or with 0.5% casamino acids for recovery of the *pheA*::Tn mutant. Colonies were counted after at least 4 weeks of incubation at 37˚C.

### Animal ethics statement

All animal protocols used in this study were reviewed and approved by the University of Minnesota Institutional Animal Care and Use Committee (IACUC) under protocol numbers 1912-37660A and 2210-40465A. The University of Minnesota’s NIH Animal Welfare Assurance Number is D16-00288 (A3456-01), expiration date 04/30/2024. All animal experiments were done in strict accordance with the recommendations in the Guide for the Care and Use of Laboratory Animals of the National Institutes of Health [83].

### Statistical analysis

A student’s unpaired t-test (two-tailed) was used for pairwise comparisons between WT, mutant and complemented strains. *P* values were calculated using GraphPad Prism 8.0 (GraphPad Software, Inc). P values < 0.05 were considered statistically significant.

### Custom scripts and code

All custom scripts and R code used for Illumina sequence read processing are available at https://github.com/bloc0078/umn-tischler-tnseq. Custom R code used for TnseqDiff analysis is provided in the supplementary material **S1 Files.**

### Data availability

All raw sequencing data from whole genome sequencing and Tn-seq experiments underlying the results reported are publicly available in FASTA format at the NCBI Sequence Read Archive (BioProject PRJNA1006392). All raw numerical and image data underlying the figures are provided in the supplementary materials.

## ACKNOWLEDGEMENTS

This work was completed in part with resources, including instrumentation and staff, at the University of Minnesota Genomics Center (UMGC; RRIC SCR_012413) and the University of Minnesota Imaging Centers (UIC: RRID SCR_020997). We thank Drs. Yusuke Minato and Anthony Baughn for developing the custom MtbYM rich media and for helpful discussions. We thank Dr. Lili Zhao for providing the updated R code for TnseqDiff analysis.

## SUPPORTING INFORMATION

**S1 Table. *M. tuberculosis* Tn mutants in M-ES genes recovered from an arrayed Tn mutant library.**

**S2 Table. Tn-seq read counts at each Tn insertion site for all experimental conditions.**

**S3 Table. Complete TnseqDiff analysis to identify Tn mutants with differential fitness across experimental conditions.** Comparisons between conditions are indicated in each tab. Bolded text indicates Tn mutants that met statistical significance cutoffs (log_2_ fold change > ± 2, adjusted *P* value <0.025.

**S4 Table. Oligonucleotide primers.**

**S1 Fig. TnseqDiff analyses to identify Tn mutants with fitness defects in *in vitro* input controls and mouse spleens.** Volcano plots of TnseqDiff statistical analyses of Tn-seq data to determine relative Tn mutant fitness in (**A)** infection input liquid culture (Liquid) vs. M-ES Tn library control (Library), **(B)** plate recovery of infection culture (Plate) vs. M-ES Tn library control (Library), **(C)** spleens at day 1 post-infection (D1 Spleen) vs. plated input control (Plate)**, (D)** spleens at day 7 vs. day 1 post-infection, and **(E)** spleens at day 42 vs. day 1 post-infection. Dashed lines indicate cutoffs for statistical significance of ± 2 log_2_ fold change and adjusted P value of <0.025. Tn mutants meeting these significance cutoffs are colored and labeled.

**S2 Fig. The *M. tuberculosis purF*::Tn mutant requires exogenous hypoxanthine and thiamine for *in vitro* growth.** Strains were grown in 7H9 supplemented with 60 μM thiamine and 150 μM hypoxanthine, washed with PBS-T, and diluted to OD_600_ = 0.05 in 7H9 broth (wild-type and *purF*::Tn), 7H9 with 150 μM hypoxanthine (*purF*::Tn +hypoxanthine), or 7H9 with 60 μM thiamine and 150 μM hypoxanthine (*purF*::Tn +hypoxanthine +thiamine). Growth was measured by optical density at 600 nm. Data represent means ± standard errors of biological triplicate cultures.

**S3 Fig. *M. tuberculosis* strains tested individually in mice are PDIM-proficient.** Thin-layer chromatographic analysis of apolar lipids extracted from cultures of WT Erdman (1) *purF*::Tn (2), *pheA*::Tn (3), *purF*::Tn pMV-*purF* (4), and *pheA*::Tn pMV-*pheA* (5) labelled with [^14^C]-propionate, which is preferentially incorporated into PDIM.

**S1 Data. Supporting data files for all graphical figures.** This supporting data file includes the individual numerical values used to generate the graphs in Fig 1, Fig 2B, Fig 2C, Fig 3, Fig 4, and S2 Fig and the results of statistical analyses on these data. Data for each figure are provided in separate tabs. Raw data used for generating the volcano plots in Figs 2D-F and S1 Fig are provided in S3 Table.

**S2 Data. Raw images.** The raw images used to generate S3 Fig are provided.

**S1 Files. Updated R code for TnseqDiff.**

