## Supplementary figures and images for "*Mycobacterium tuberculosis* requires conditionally essential metabolic pathways for infection"

### Supplemental Figures

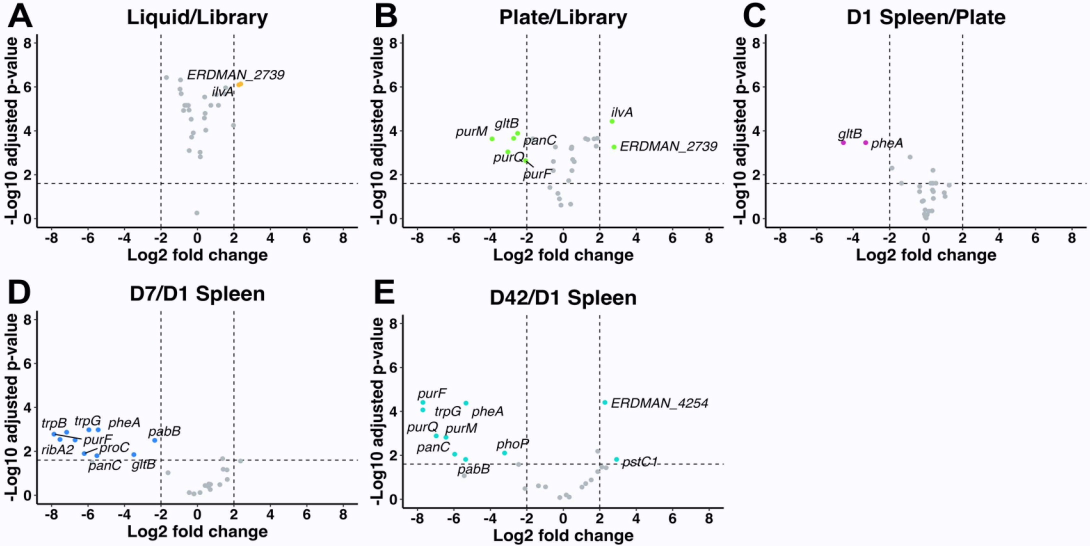

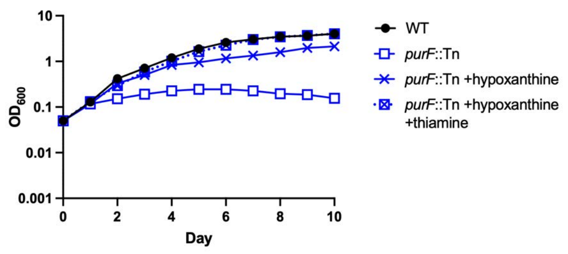

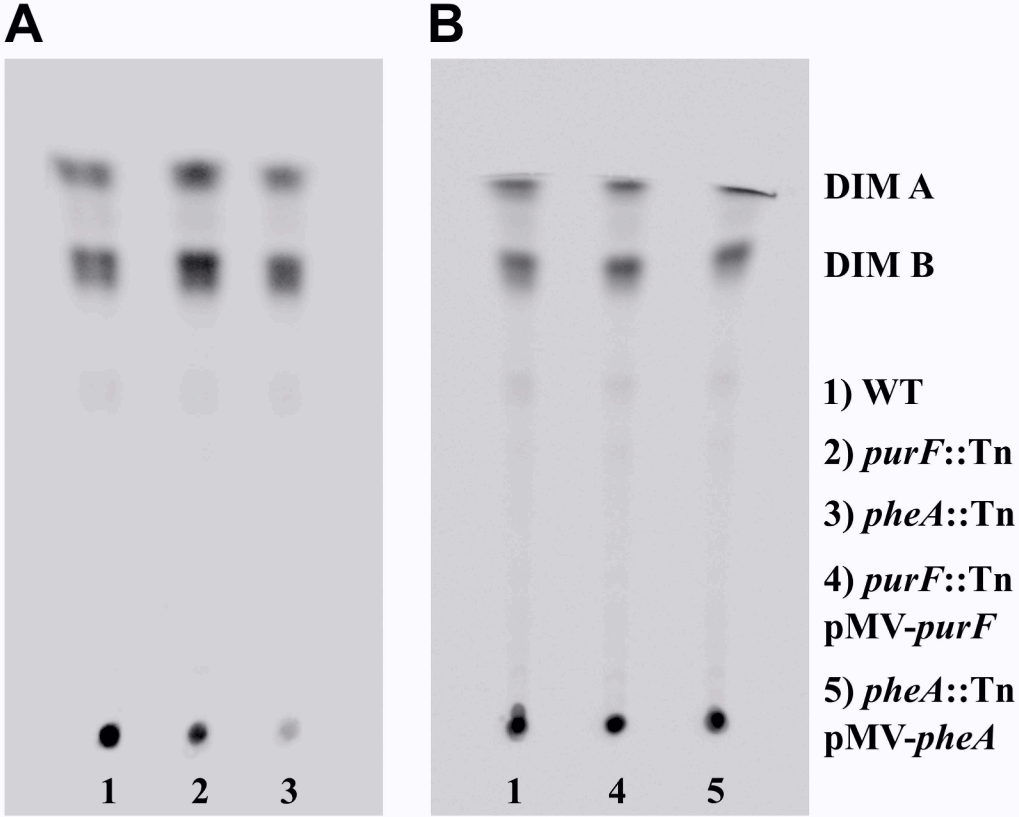
